# A comprehensive urine proteome database generated from patients with various renal conditions and prostate cancer

**DOI:** 10.1101/2021.02.10.430660

**Authors:** Adam C. Swensen, Jingtang He, Alexander C. Fang, Yinyin Ye, Carrie D. Nicora, Tujin Shi, Alvin Y. Liu, Tara K. Sigdel, Minnie M. Sarwal, Wei-Jun Qian

## Abstract

Urine proteins can serve as viable biomarkers for diagnosing and monitoring various diseases. A comprehensive urine proteome database, generated from a variety of urine samples with different disease conditions, can serve as a reference resource for facilitating discovery of potential urine protein biomarkers. Herein, we present a urine proteome database generated from multiple datasets using 2D LC-MS/MS proteome profiling of urine samples from healthy individuals (HI), renal transplant patients with acute rejection (AR) and stable graft (STA), patients with non-specific proteinuria (NS), and patients with prostate cancer (PC). A total of ~28,000 unique peptides spanning ~2,200 unique proteins were identified with a false discovery rate of <0.5% at the protein level. Over one third of the annotated proteins were plasma membrane proteins and another one third were extracellular proteins according to gene ontology analysis. Ingenuity Pathway Analysis of these proteins revealed 349 potential biomarkers. Surprisingly, 43% (167) of all known cluster of differentiation (CD) proteins were identified in the various human urine samples. Interestingly, following comparisons with five recently published urine proteome profiling studies, which applied similar approaches, there are still ~400 proteins which are unique to this current study. These may represent potential disease-associated proteins. Among them, several proteins such as myoglobin, serpin B3, renin receptor, and periostin have been reported as pathological markers for renal failure and prostate cancer, respectively. Taken together, our data should provide valuable information for future discovery and validation studies of urine protein biomarkers for various diseases.

## Introduction

The production and elimination of urine is essential for the removal of waste products generated by cellular metabolism and other processes. Kidneys use special structures, particularly glomeruli, to filter blood.^1, 2^ Important substances such as water, salts, glucose, other nutrients, and most proteins are reabsorbed by the kidneys. Only select proteins are removed for excretion in urine. Therefore, urine protein excretion in healthy adults is usually limited to less than 150 mg/day.^3^ Urine protein excretion beyond this value is defined as proteinuria,^4^ which is often a sign of kidney damage. The proteins in urine can originate from the kidney, bladder, prostate gland, ureter, urethra, or even from distant organs and tissues. Since urine can be collected in large quantities using non-invasive procedures, urine proteins are particularly suitable for use as biomarkers to diagnose and monitor dysfunction involving these organs. Some urine protein biomarkers are critical for diagnosing and monitoring diseases such as prostate cancer^5, 6^ and kidney failure.^7–9^ To facilitate the discovery of novel urine protein biomarkers, it is necessary to generate a comprehensive urine protein database from samples collected from patients with various disease conditions and healthy patients.

Mass spectrometry (MS)-based proteomics provides a powerful analytical tool for large-scale identification of proteins found in urine. There have been many urine proteome profiling studies using different separation approaches coupled with MS. For instance, Adachi et al. employed SDS-PAGE and reverse phase liquid chromatography (LC) for protein separation and identified a total of 1,543 proteins from urine samples of healthy individuals via LC-MS/MS analysis using LTQ-FT and LTQ-Orbitrap mass spectrometers.^10^ Using SDS-PAGE and lectin enrichment followed by LC-MS/MS, Marimuthu et al. identified 1,823 proteins from healthy human urine.^11^ Gel-free methods have also been used for urine proteome profiling. For example, Li et al. applied a multidimensional LC-MS/MS method and identified 1,310 urine proteins.^12^ Expanded coverage of 3000-6000 proteins from the human urine proteome have been recently reported by applying more complex ligand library bead-binding equalization techniques or multi-dimensional gel electrophoresis coupled with multi-dimensional LC-MS/MS approaches^13, 14^. However, one of the major limitations of these urine proteome profiling studies was the focus on only healthy individuals such that many disease-associated proteins could be missed from these studies. Therefore, it would be valuable to have a comprehensive urine proteome database derived from both healthy and disease conditions as a reference resource for guiding urine protein biomarker discovery.

In our previous studies, we have performed comparative studies of urine of renal patients and healthy individuals with the purpose of identifying potential urinary protein biomarkers for acute renal transplant rejection.^15, 16^ In order to generate a urine proteome database originating from multiple disease conditions as a reference resource, we combined datasets from urine samples from patients suffering from prostate cancer, renal transplant, and non-specific proteinuria, as well as healthy individuals using a commonly applied 2D-LC-MS/MS workflow. Urine proteins in each group of samples were digested into peptides which were pre-fractionated by either strong cation exchange or high-pH reversed-phase LC. Peptides in each fraction were analyzed by LC-MS/MS, resulting in the identification of a total of ~28,000 unique peptides across ~2,200 urinary proteins. The final database was annotated with observation counts from each biological condition as well as the annotation of presence or absence in five recent urine proteome profiling studies. Approximately 400 proteins were only observed in the current study, possibly suggesting the observation of disease-associated proteins. Since the database was generated from several disease conditions and annotated against other urine proteome databases from healthy individuals, our database could serve as a global reference for future biomarker discovery studies using urine as the source sample.

## 2. Materials and Methods

### 2.1 Urine collection and processing

A total of 45 urine samples from 10 renal transplant patients with proven acute rejection (AR), 10 renal transplant patients with stable graft (STA), 10 non-specific proteinuria patients (NS), 10 healthy individuals (HI), and 5 prostate cancer (PC) patients were utilized for global urine proteome profiling. The patient demographics of patients with renal conditions including healthy controls were the same as described previously with an age range of 3 to 21^15^. The PC urine samples were from pre-operation patients with an age range of 60 to 75. This research was approved by the Institutional Review boards at Stanford University, University of California San Francisco, University of Washington, and Pacific Northwest National Laboratory in accordance with federal regulations. ~50 mL urine samples were collected from each patient in sterile containers. Samples were centrifuged at 2000 × g for 20 min at room temperature within 1 h of collection. The supernatant was collected and stored at −80 °C for further analysis.

Proteins in the urine supernatant were concentrated with 10-kDa Amicon Ultra-15 centrifugal filter units (Millipore). The final protein concentration was measured by bicinchoninic (BCA) assay (Pierce). After concentration, 45 μg of urine proteins were pulled from each sample and combined according to their clinical categories, namely, PC, AR, STA, NS and HI. The pooled protein samples were denatured by 8 M urea, reduced by 10 mM dithiothreitol (DTT), alkylated by with 40 mM iodoacetamide, and digested by trypsin as previously described.^15^ The final peptide concentrations were measured using the BCA assay.

Peptides from pooled AR, pooled STA, pooled NS, and pooled HI samples were fractionated by strong cation exchange (SCX) chromatography as previously described[17], and peptides from pooled PC samples were fractionated by high-pH reversed-phase separation and concatenated into 24 fraction as previously described.^17^

### 2.2 LC-MS/MS analyses

The peptide fractions were analyzed by LC-MS/MS. Specifically, the peptide fractions from SCX were analyzed by LTQ linear ion trap mass spectrometer (Thermo Fisher Scientific) coupled with a customized LC system as previously described.^15^ The peptide fractions from high pH reversed-phase LC fractionation were analyzed an LTQ Orbitrap Velos mass spectrometer (Thermo Fisher Scientific) coupled with a similar customized LC system. LC columns were prepared in-house by slurry packing 3-μm Jupiter C18 (Phenomenex, Torrence, CA) into 35-cm × 75μm i.d fused silica (Polymicro Technologies Inc., Phoenix, AZ). A 100-min LC gradient with a 300 nL/min flowrate was applied for separations. The resolution of the MS scan was 120,000 with Top-20 data-dependent MS/MS acquisitions on CID mode.

### 2.3 Proteomics data analysis

All MS/MS spectra were searched against the UniProtKB/Swiss-Prot protein knowledgebase release 2013_09 using MSGF+ (Release 2019.07.03). The search parameters were as follows: (1) fixed modification, carbamidomethyl of C; (2) variable modification, oxidation of M; (3) allowing two missed cleavages; (4) parent ion mass tolerance: 1.0 Da for LTQ data and 20 ppm for Orbitrap Velos data; (5) fragment ion mass tolerance, 1.0 Da. MS Generating-Function (MSGF) scores were generated for all identified spectra by computing rigorous p-values (MSGF SpecEvalue).^18^ The FDRs for final peptide and protein identifications were controlled to be < 0.1% and < 0.5%, respectively. Gene Ontology (GO) annotation for cellular component and biological process of the identified urine proteins was performed by using the Database for Annotation, Visualization and Integrated Discovery (DAVID 6.8) bioinformatics resource.^19, 20^ Biomarkers were screened from the identified proteins using the Ingenuity Pathway Analysis (IPA Ver. 48207413) biomarker filter module.

#### Data availability

The datasets presented in this study can be found in online repositories. The names of the repository/repositories and accession number(s) can be found below: Massive.ucsd.edu with accession: MSV000086484.

## 3. Results and Discussion

### 3.1 Global profiling of the urine proteome

Since many of the previous urinary proteomics studies reported on healthy subjects, our purpose here was to create a urine proteome database with urinary proteins that would be detectable in both disease and healthy conditions. Importantly, we wanted to demonstrate what could be detected using commonly applied, translatable, LC-MS/MS techniques. The concept was to combine datasets from samples related to renal or other conditions relevant to the urinary tract from several global urine proteome profiling efforts from our laboratory as summarized in **Figure 1,** including an illustration of potential sources of urinary protein biomarkers from different disease conditions. The first profiling efforts involved pooled urine samples from a renal rejection study with four different clinical conditions (AR, STA, NS, HI); where following protein digestion, peptides were fractionated into 32 fractions per sample by strong cation exchange chromatography (SCX) and analyzed by LC-MS/MS on LTQ. The second study involved pooled samples from prostate cancer patients; where peptides were fractionated by high-pH reversed-phase LC into 24 fractions and analyzed by LC-MS/MS on Orbitrap Velos. The combined dataset includes a total of ~150 LC-MS/MS analyses to generate the final urine proteome database. Following database searching with the MSGF+ algorithm, we identified a total of ~28,000 unique peptides (**Table S1**) and ~2,200 unique proteins (**Table S2**) with FDR <0.5% at the protein level based on decoy database search and using stringent filtering criteria.

**Figure 1.**
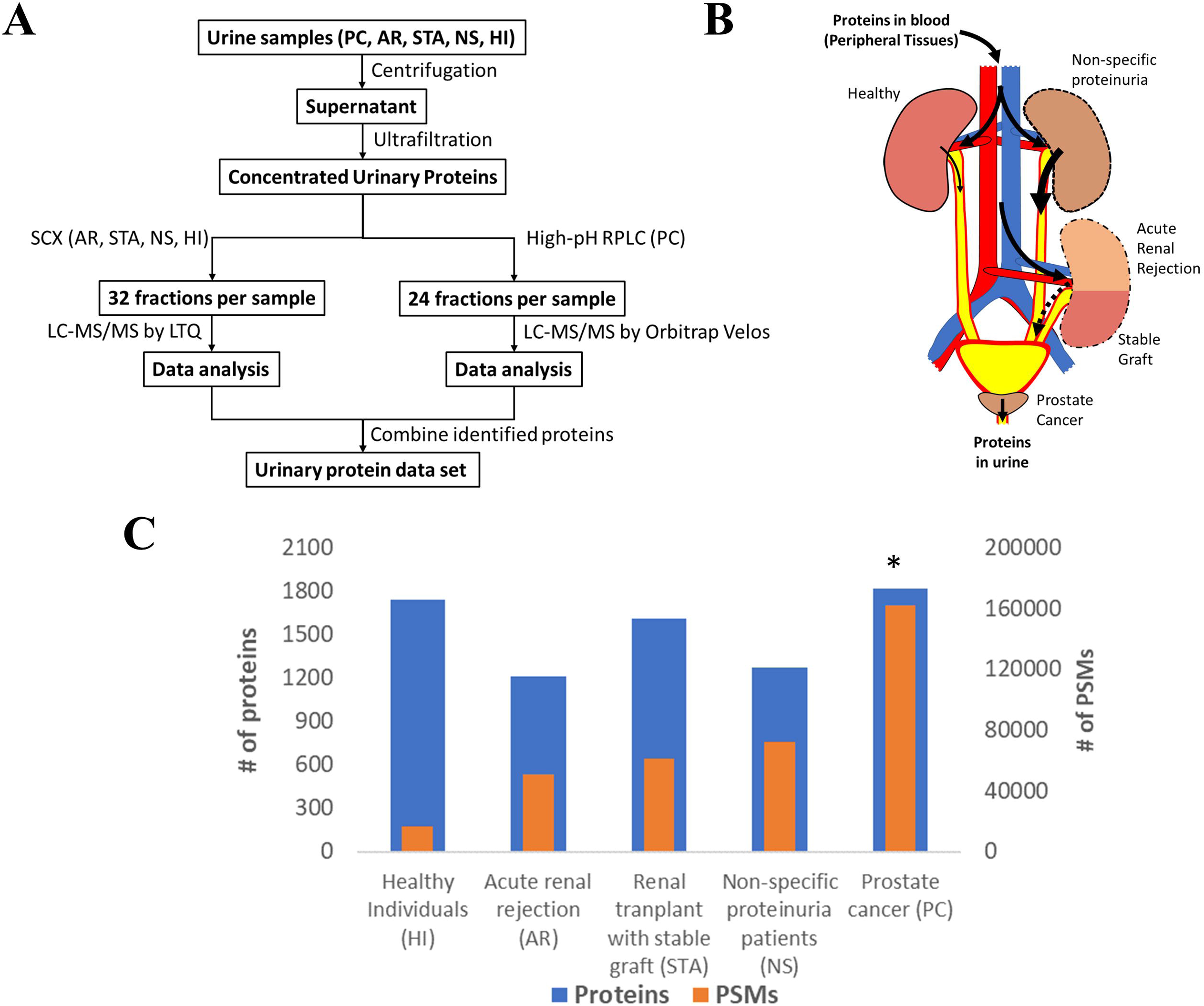
**A)** An overview of the workflow for analysis of the urine proteome; **B)** An illustration of potential sources of protein biomarkers from different organs into urine; C) The relative number of proteins and PSMs across conditions.

**Figure 1C** shows the number of total proteins and total PSMs from different conditions. Although two different sets of methods were used in this study, the proteome coverage in terms of the number of proteins identified from each condition was still relatively comparable. However, the count of PSMs in all disease conditions were substantially higher than the healthy condition, suggesting the potential leakage of highly abundant proteins into urine in these disease conditions (**Figure 1C**). For example, we observed 27,474 and 18,116 PSMs for serum albumin in the NS and AR conditions, respectively, but only 603 PSMs for albumin in the HI condition. Several other highly abundant serum proteins such as serotransferrin, retinol-binding protein 4, protein AMBP, and alpha-1 antitrypsin were also observed with high PSMs in disease conditions compared to HI.

### 3.2 Comparison to previous urine profiling or biomarker studies

Next, we compared our current set of urine proteins with results from several recent published global urine proteome profiling studies, which were mostly from healthy donors. Using a combination of 15 prior studies (including most listed in **Figure 2**), Farrah et al. compiled a comprehensive list for the PeptideAtlas project urine section (UrinePA)^21^ and found 2491 unique proteins confidently detected in those studies. The current most accessible proteome profiling method is the 2D-LC-MS/MS workflow, which consists of pre-fractionation with either high pH reverse phase LC or strong cation exchange (SCX) chromatography. Several groups have used this method to obtain relatively high urine proteome coverage.^12, 22^ Other specialized techniques have also been applied to expand the urine proteome coverage, including micro-vesicle and exosome enrichment prior to MS sample preparation.^13^, 2D SDS-PAGE separation and spot excision followed by LC-MS^10, 11, 23^, multi-dimensional gel electrophoresis followed by multi-dimensional LC-MS^14^, and using combinatorial peptide ligand library (CPLL) binding beads to “equalize” protein abundances^22^. A urine proteome database with deeper coverage (~6,000 proteins) was recently reported using several specialized methods^14^. However, it is still unclear whether disease associated proteins would be missed from these efforts focusing on samples from healthy individuals. Our urinary proteome database was generated using easily accessible techniques and incorporated multiple disease conditions in order to provide a useful baseline reference resource for guiding urine biomarker discovery. The comparison of the urine proteome coverage from different studies using various techniques is illustrated in **Figure 2**.

**Figure 2.**
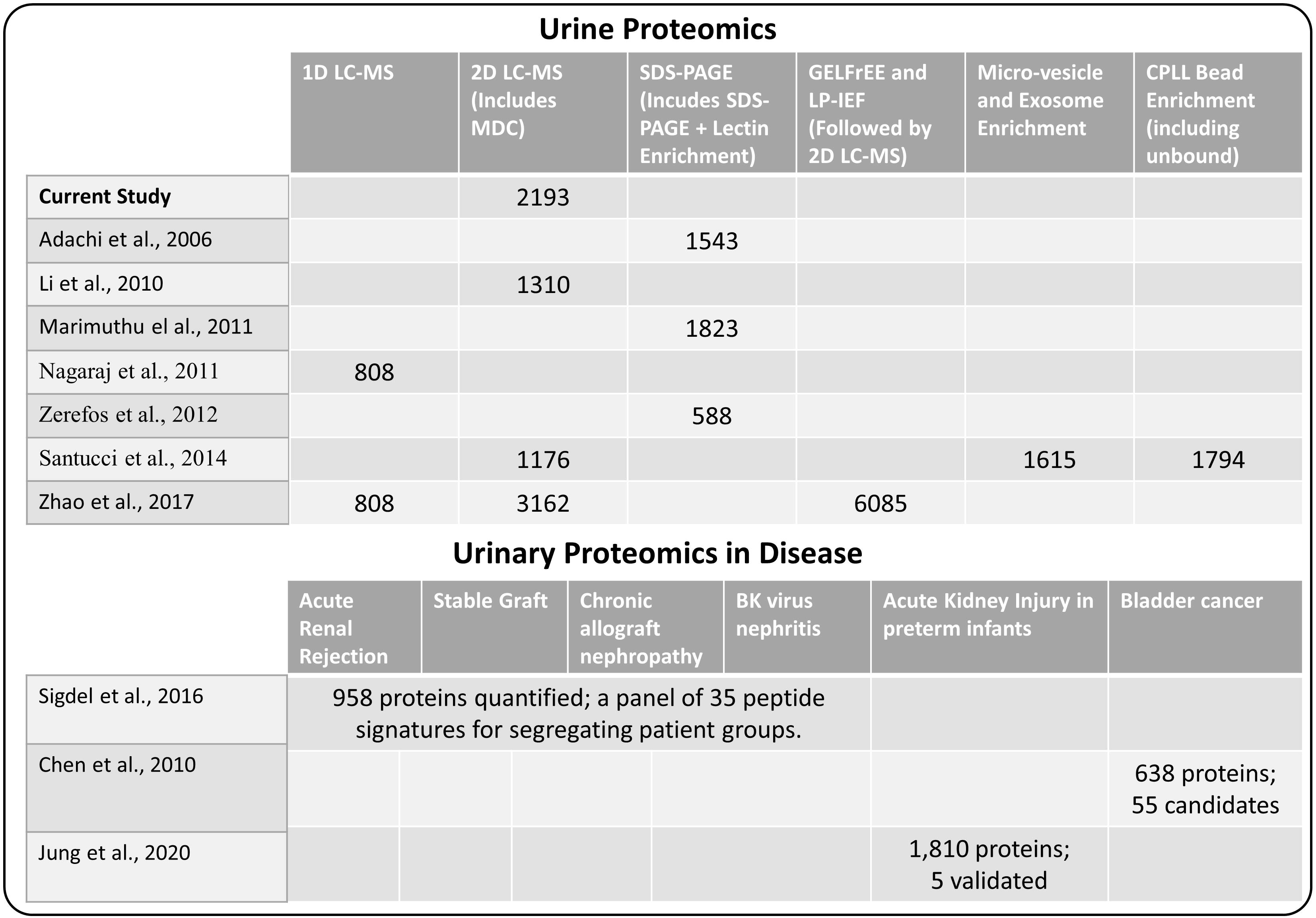
**A)** Comparison of methods used for urine proteome analysis and the number of proteins detected using the various methods **B)** Highlights of several recent biomarker discovery and verification studies in several disease conditions.

One of the primary interests of urine proteomics is the discovery of novel biomarkers of various diseases^24^. While most urine biomarker studies were based on validating target panel of protein or peptide signatures in different disease conditions, there were several studies applying global discovery approach followed by targeted verification of candidate markers. We also highlighted several biomarker discovery and verification efforts in **Figure 2**, including the study by Sigdel et al. for renal transplantation conditions^16^, the bladder cancer study by Chen et al.^25^, and the acute kidney injury study in preterm infants by Jung et al.^26^ Importantly, all of the reported candidate biomarkers from these studies were identified in our current dataset (**Table S2**), again suggesting the relative comprehensiveness of the current urine proteome database.

### 3.3 Gene ontology analysis of the urine proteome

The urine proteins identified in this work were classified based on Gene Ontology (GO) cellular component and biological process annotation terms using the Database for Annotation, Visualization and Integrated Discovery (DAVID 6.8) bioinformatics resources. Of note, urine proteins were annotated as extracellular space and plasma membrane proteins at 27.4% and 43.3%, respectively (**Figure 3A**). Adachi et al. and Marimuthu et al. also found that plasma membrane proteins were enriched in urine samples, where ~ 20% ^10^ and 31% ^11^ of urine proteins identified from healthy human urine were plasma membrane proteins. In terms of biological processes, 24.4% proteins were found to function in cell adhesion, which is consistent with the previously mentioned enrichment of extracellular space and plasma membrane proteins. It is well known that the key protein components involved in various cell-adhesion structures (adherens junction, focal adhesion, desmosome, tight junction, and so on) are localized in the plasma membrane and extracellular matrix. Many examples of these proteins (cadherins, desmocollins, desmogleins, integrins, collagens and fibronectins) were identified in this work. It is not surprising that cell-adhesion related proteins are enriched in urine because they are exposed on cell surfaces which increases the likelihood of release into urine. Besides cell adhesion, there are several other major biological processes that are enriched, including proteolysis (19.7%), immune response (18.5%), cell proliferation (16.4%), and response to wounding (9.7%) (**Figure 3B**).

**Figure 3.**
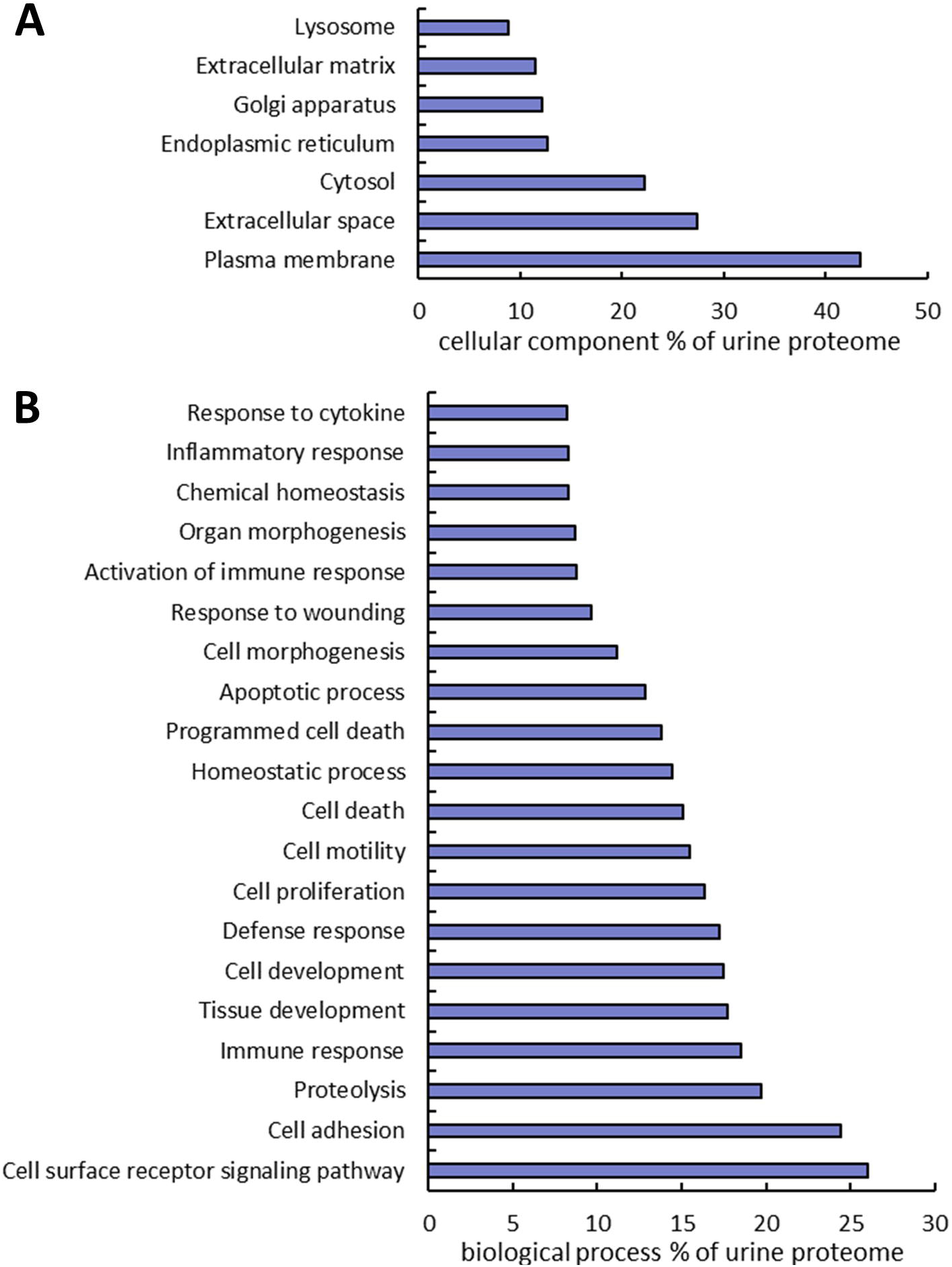
Gene Ontology annotation of identified proteins as a percent of the urine proteome. GO cellular component **(A)** and biological process **(B)** terms were derived using the DAVID bioinformatics database.

### 3.4 Cluster of Differentiation (CD) antigens

An important feature of our dataset is that many CD antigens were identified from human urine. CD antigens often play critical roles in cell signaling and cell adhesion. They are commonly used as markers for immunophenotyping and for diagnosis, monitoring, and treatment of diseases. Out of all 394 known human CD proteins in the Uniprot database, 178 (45%) were identified from the human urine samples in this study (**Table S4**). Since CD proteins are cell-surface proteins, it is not a surprise that many CD molecules are released from cells into body fluids such as blood and urine. Also worthy of note, two extensively used prostate cancer stem cell markers CD133 ^33^ and CD44 ^34^ were identified. These CD proteins could be useful as diagnostic markers for other diseases.

### 3.5 Candidate biomarkers identified from the urine proteome

Due to the non-invasive nature of urine collection, urine proteins are ideal biomarkers for diagnosis of renal diseases and other diseases related to the urinary tract including prostate and bladder cancer. Indeed, several promising biomarkers have already been reported.^27–32^ Herein we performed a “biomarker filter” analysis of the identified urine proteins using the Ingenuity Pathway Analysis (IPA Ver. 48207413) software. The “biomarker filter” is an IPA module which allows identification of biomarker candidates based on prior curated literature data. 349 proteins were identified as candidate biomarkers (**Table S5**) following biomarker filtering analysis. These biomarkers were categorized based on their applications. 153, 108, 71, and 35 proteins were relevant to diagnosis, efficacy, prognosis, and disease progression, respectively (**Figure 4B**). We also analyzed the tissue specificity of 349 tissue-enriched proteins based on human protein atlas database (**Figure 4A**). Of these, 230 were expressed in kidneys, 200 were expressed in the prostate gland, and 181 were expressed in bladder, among many other represented tissues (**Table S6)**. However, most of these proteins were shared by multiple tissues or organs.

**Figure 4.**
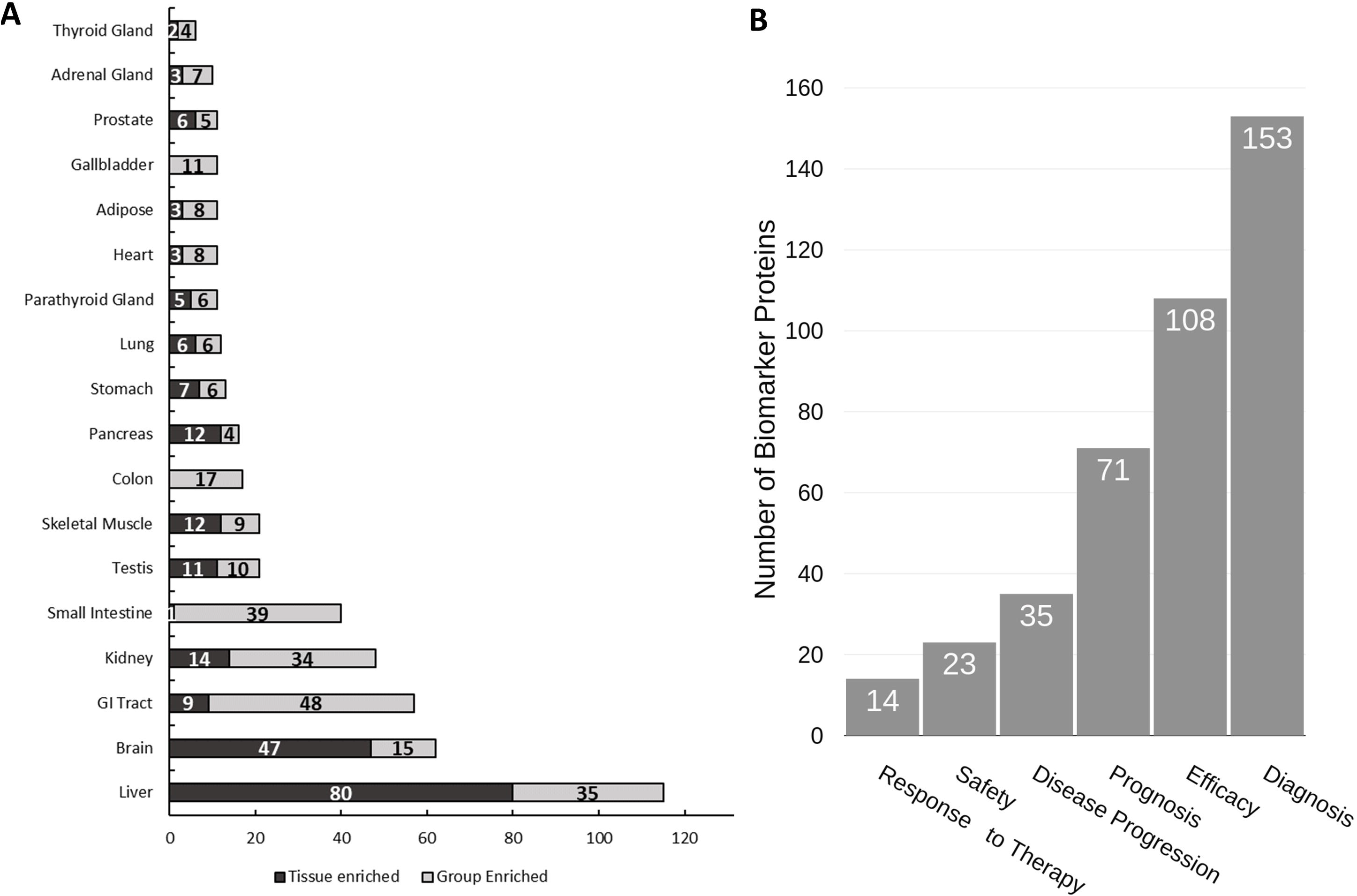
Analysis of urine proteome for tissue specificity and disease biomarkers. **A)** Tissue specificity of the urine proteome was derived from the Human Protein Atlas database (https://www.proteinatlas.org). **B)** Functional utility of detected disease biomarkers found in urine as annotated by IPA. Note that tissue enrichment was defined by the Human Protein Atlas to be expression in a single tissue at least five-fold greater than that of all other tissues. Group enrichment was defined by the Human Protein Atlas to be a five-fold greater average expression level in a group of two to seven tissues compared to all other tissues.

### 3.6 Potential disease-associated proteins

Another potential value of our dataset is to identify potential disease-associated proteins by comparing to the multiple previously published datasets based on samples from healthy individuals. We were able to compare our data to five existing datasets where comparable profiling approaches were applied^10–14^; differing primarily by the extensiveness and complexity of pre-fractionation utilized. Indeed, ~400 unique proteins were only observed our dataset (**Table S3**). We note that while some of the unique proteins may be due to protein accession ID discrepancies between different studies, the data suggest the detection of disease-associated proteins. As shown in **Table 1**, a number of histone proteins were predominantly detected in prostate cancer samples, supporting the recent report that extracellular or circulatory histones may reflect tissue injury or cell death.^33^ The detection of much higher abundant myoglobin in renal conditions only in our dataset is an interesting confirmation since urine myoglobin was often used as a marker for acute renal failure.^34^ Moreover, Serpin B3 was detected at highest level in the renal transplant patients with stable graft (STA), consistent with the report that Serpin B3 was an important healing biomarker. On the other hand, detection of high level of insulin in the prostate cancer samples is also worthy of noting since the older individuals are much likely to have higher levels of circulating insulin due to insulin resistance.^35^ Renin receptor and Periostin were two markers detected primarily in prostate urine samples and both proteins have been reported as viable cancer biomarkers^36,37^.

**Table 1.**
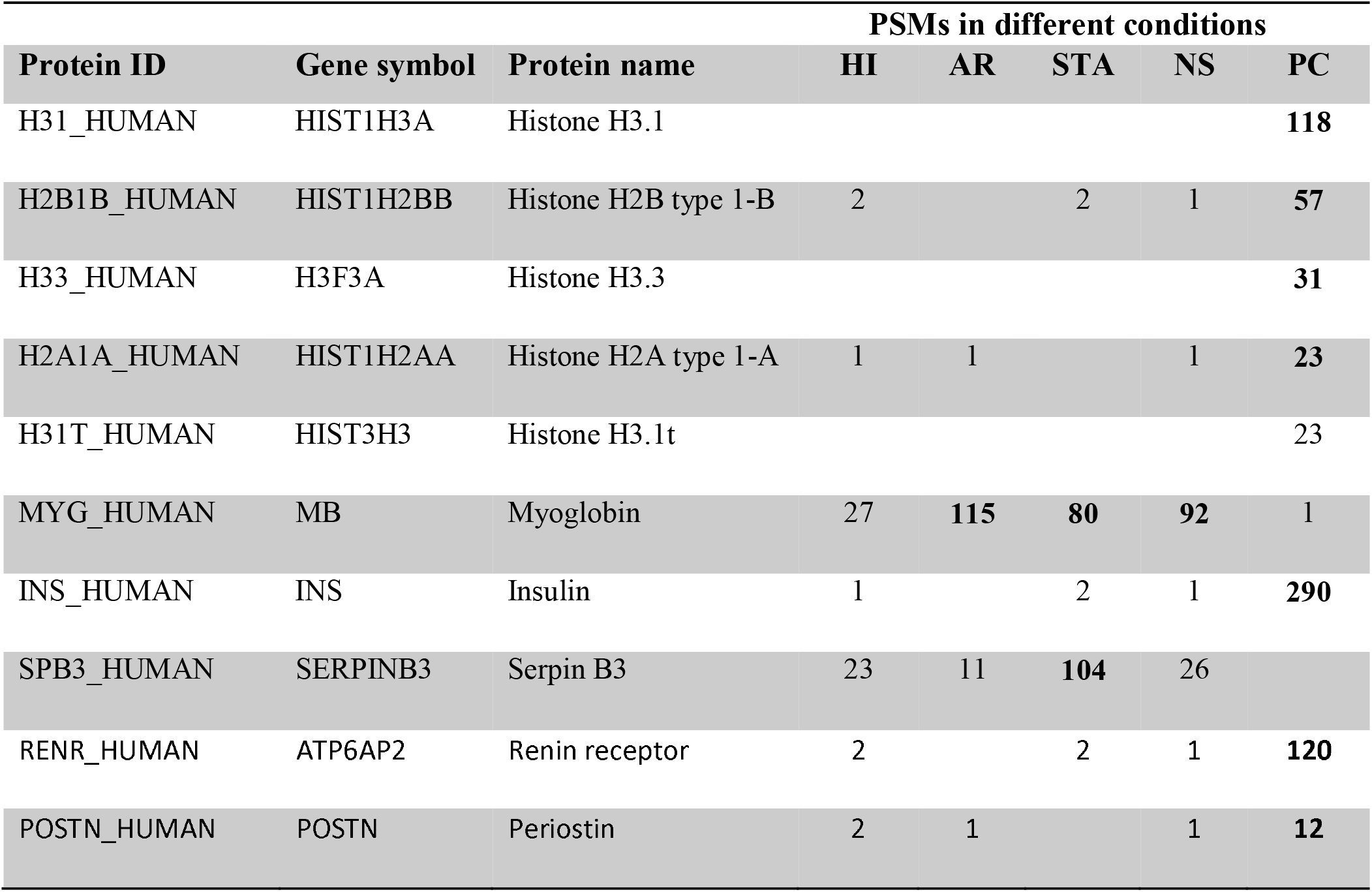
Selected potential disease-associated proteins only detected in the current study.

### 4. Concluding Remarks

We have generated a comprehensive urine proteome database through LC-MS/MS profiling of urine samples from prostate cancer patients, renal transplant patients with acute rejection or stable graft, non-specific proteinuria patients, and healthy individuals. The overall analyses resulted in the identification of ~28,000 unique peptides and ~2,200 unique proteins. Over 40% of the identified proteins were annotated as plasma membrane proteins and over three-fourths were extracellular proteins. IPA biomarker filter analysis revealed that 349 proteins are potential candidate biomarkers relevant to diagnosis, efficacy, prognosis, and disease progression. Moreover, 45% (178) of all known CD proteins were identified in these human urine samples. Presumably due to the inclusion of several disease conditions, our study identified ~400 proteins that were not detected in previous profiling studies using similar approaches. Among them, several interesting disease-associated protein markers were identified. Together, this comprehensive urine proteome dataset could serve as a valuable reference resource for future biomarker discovery efforts using urine as the source sample.

## Abbreviations

AR: acute rejection
CD: Cluster of Differentiation
DAVID: Database for Annotation, Visualization and Integrated Discovery
GO: Gene Ontology
HI: healthy individuals
IPA: Ingenuity Pathway Analysis
LC: Liquid Chromatography
MS: Mass Spectrometry
MSGF: MS Generating-Function
NS: non-specific proteinuria
STA: stable graft
PC: prostate cancer
PSM: Peptide-Spectrum Match

## Acknowledgements

Portions of the research were supported by the DoD PCRP Program Award W81XWH-16-1-0614. The proteomics experimental work described herein was performed in the Environmental Molecular Sciences Laboratory, PNNL, a national scientific user facility sponsored by the DOE under Contract DE-AC05-76RL0 1830.

## Author contributions

AYL, TKS, MMS, and WJQ designed the experiments, CDN and TS performed the experiments, ACS, JH, ACF, and YY analyzed the data, ACS, JH, ACF, and WJQ wrote the manuscript.

## Supplemental Tables

**Supplemental Table S1**. A list of all peptide spectral matches (PSMs) that were detected in our combined dataset; and the count of spectra wherein they were detected (spectral count). Peptide were identified using MSGF+. The confidence score is included (lower is better). Sequences include preceding and training amino acids of the protein sequence not part of the detected sequence (i.e., sequences are contained within the periods/full stops “.”).

**Supplemental Table S2**. A list of all the proteins from which peptides were detected in our combined dataset. The number of unique PSMs and the sum-totaled spectral counts for the corresponding PSMs are included. The most confident MSGF_SpecProb peptide score is included (lower is better). The presence of the proteins in previous urine proteome profiling studies or biomarker studies was also annotated.

**Supplemental Table S3**. List of proteins that are not detected by previous five urine proteome studies using 2D LC or comparable approaches

**Supplemental Table S4.** A list of Cluster of Differentiation (CD) proteins detected in urine in this study.

**Supplemental Table S5**. Ingenuity Pathway Analysis (IPA) generated list of proteins known to be useful as biomarkers. Where known, the cellular location and protein family are included. The type of biomarker (e.g., diagnosis, efficacy, prognosis, etc.) and the diseases for which the biomarker is useful are included.

**Supplemental Table S6**. A list of all the proteins detected in our dataset and their corresponding tissue and cell-type specificities. Specificities were determined based on detectability from the Human Protein Atlas (Uhlen et al. “https://www.proteinatlas.org”). Cell location, protein family, and known targeting drugs are included where available.

